# Identification of peptides interfering the lrrk2/pp1 interaction

**DOI:** 10.1101/807487

**Authors:** Chang Zhi Dong, Heriberto Bruzzoni-Giovanell, Yanhua Yu, Karim Dorgham, Christophe Parizot, Jean Marc Zini, Pierre Tuffery, Angelita Rebollo

## Abstract

Serine/threonine phosphatases are responsible for counteracting the effect of the protein kinases implicated in the development of several pathologies. Here we identified by PEP-scan approach the sequence of a fragment of LRRK2, a Parkinson’s disease associated protein, interacting with the phosphatase PP1. The fragment, that is located in a LRRK2 domain of undefined function, was associated in N-terminal to an optimized cell penetrating peptide in order to study their in vitro and *in vivo* biological activity. From this original sequence, we developed and studied five interfering peptides (IPs) and identified two peptides able to disrupt the LRRK2/PP1 interaction by *in vitro* competition in anti-LRRK2 immunoprecipitates. Using FITC-labelled peptides, we confirmed the internalization of the peptides in cell lines as well as in and primary human normal and pathological cells. Finally, we have confirmed by ELISA test the association of Mut3DPT-LRRK2-Long and Mut3DPT-LRRK2-Short peptides to purified PP1 protein in a selective manner. The shortest peptides, MuteDPT-LRRK2-5 to 8 with either N or C-terminal deletions are not able neither disrupt the association LRRK2/PP1 nor to associate to purified PP1 protein. The peptides Mut3DPT-LRRK2-Long and Mut3DPT-LRRK2-Short may be new tools to study the role of LRRK2/PP1 interaction in normal and pathological conditions.

## 1 INTRODUCTION

Serine/threonine protein phosphatases 1 and 2A (PP1 and PP2A) are the most ubiquitous and abundant serine/threonine phosphatases in eukaryotic cells and they are implicated in the regulation of various essential cellular functions ^1,2^. PP1 is a single-domain hub protein with nearly 200 validated interactors in vertebrates. It forms stable complexes with PP1-interacting proteins (PIPs) that guide the phosphatase throughout its life cycle and control its fate and function. The binding of PP1 to PIPs is mainly mediated by short motifs that dock to surface grooves of PP1; most of them combine short linear motifs to form large and unique interaction interfaces with PP1 ^3,4^. Although PIPs often contain variants of the same PP1 binding motifs, they differ in the number and combination of docking sites. This combinatory strategy for binding to PP1 creates holoenzymes with unique properties. PIPs control associated PP1 by blocking their interaction with other PIP or blocking the access to the active site. In addition, some PIPs have a subcellular targeting domain that promotes dephosphorylation by increasing the local concentration of PP1. Many PIPs have separate domains for PP1 anchoring, PP1 regulation, substrate recruitment and subcellular targeting, which enable them to direct associated PP1 to a specific subset of substrates and mediate acute activity control. Hence, PP1 functions as the catalytic subunit of a large number of multimeric holoenzymes, each with its own subset of substrates and mechanism(s) of regulation. The diversity of the PP1 interactome and the properties of the PP1 binding code account for the exquisite specificity of PP1 *in vivo* ^5,6^.

Many naturally occurring protein phosphatase inhibitors with different relative PP1/PP2A affinities have been described and are widely used as powerful research tools. In particular microcystins are highly toxic cyanotoxins by binding to their main targets PP1 and PP2A ^6^. In the same direction, okadaic acid has been used to specifically inhibit PP1 or PP2A, depending on the used concentration ^7^.

Leucine-Rich Repeat Kinase 2 (LRRK2) has been identified as a PP1 interacting protein ^8^. Mutations in the LRRK2 gene are a common cause of familial Parkinson’s disease (PD). Variation around the LRRK2 locus also contributes to the risk of sporadic PD although the involved mechanism is non-well understood. A cluster of phosphorylation sites in LRRK2 ^9-11^, including Ser910, Ser935, Ser955 and Ser973, is important for PD pathogenesis as several PD-linked LRRK2 mutants are dephosphorylated at these sites. LRRK2 is also dephosphorylated in cells after pharmacological inhibition of its kinase activity, which is currently proposed as a strategy for disease-modifying PD therapy. PP1 is responsible also for LRRK2 dephosphorylation observed in PD mutant LRRK2 and after LRRK2 kinase inhibition ^8,10^.

Protein phosphatases have both protective and promoting roles in the etiology of diseases. A prominent example is the existence of oncogenic as well as tumor-suppressing protein phosphatases. Several works have point out the importance of identifying phosphatase modulators ^7,12^. A few numbers of modulators of protein phosphatase activity are already applied in therapies. They were however not developed in target-directed approaches. For this reason, it will be interesting to develop targeted modulators of phosphatases, either targeting their enzymatic site or by targeting the sites of interaction with their partners (particularly those involved in pathological process) as research tools. We and others have developed interfering peptides targeting PP2A interactions and showed their potential interest in the development of therapeutic strategies ^13-15^. In the same way, we introduce now interfering peptides targeting PP1/LRRK2 interaction in order to provide new tools to understand this interaction and their biological signification, as well as their potential use as therapeutic peptide.

## 2 MATERIALS AND METHODS

### 2.1 Peptides synthesis and sequence

Fmoc/*t*Bu strategy was chosen for the peptide synthesis except for the first C-terminal residue, lysine of which the side chain was protected with dde group. Upon completion of the synthesis, the peptide-anchored resin was handled with 2% monohydrate hydrazine in DMF according to the manufacture recommended method to remove the dde group. The peptide-anchored resin was then shaked in DMF with either FITC-NCS alone (1.5 eq.) or biotin (3 eq.) in the presence of DCC (3 eq.)/HOBt (3 eq.)/DIEA (5 eq.) overnight at room temperature. After washing 4 times with DMF and 4 times with CH_2_Cl_2_, the peptide was finally cleaved from the resin and precipitated twice with cold ether/heptane (1/1). It was then dissolved in 30% CH_3_CN in water and lyophilized. The purification was performed by RP-HPLC using an increasing CH_3_CN gradient. Its identity was confirmed by maldi-mass spectrometry.

### 2.2 Peptide structure modelling

The structure of the long sequence was predicted using PEP-FOLD3 ^16^ in house implementation and subject to visual inspection to identify candidate N- and C-terminal amino acids deletions likely to question peptide interfering ability.

### 2.3 Cell lines

Human cancer breast cell line MDA-MB231 was cultured in DMEM medium supplemented with 10% foetal calf serum (FCS). Peripheral blood mononuclear cells (PBMC) were cultured in RPMI medium supplemented with 10% of FCS.

### 2.4 PP1 binding assay on cellulose-bound peptides containing LRRK2 sequence (PEP-scan)

Overlapping dodecapeptides with two amino acid shift, spanning the complete LRRK2 sequence were prepared by automatic spot synthesis (Abimed, Langerfeld, Germany) onto an amino-derived cellulose membrane, as described ^17,18^. The membrane was saturated using 3% non-fat dry milk/3% BSA (2h room temperature), incubated with purified PP1 protein (4 μg/ml, 4°C, overnight) and after several washing steps, incubated with polyclonal anti-PP1 antibody 2h at room temperature, followed by HRP-conjugated secondary antibody for 1h at room temperature. Positive spots were visualized using the ECL system.

### 2.5 Comparative modelling of the LRRK2 domain

The comparative modelling of the LRRK2 domain was done using the hhsearch suite ^19^ to identify a possible 3D template of the Protein Data Bank ^20^ 3D modelling was performed using as template the 3DPT PDB entry and the Tito software ^21^ to refine the hhsearch alignment, and identify preserved regions of the template. Loops were then built using the DaReUS-loop approach ^22^.

### 2.6 Isolation and culture of primary cells

Fresh blood from healthy donors (HD) was obtained from Etablissement Francais du Sang. Chronic lymphocytic leukemia (CLL) patient samples were obtained from the Department of Hematology. Peripheral blood mononuclear cells (PBMC) from HD and CLL patients were prepared by Ficoll gradient centrifugation. Cells were maintained in RPMI 1640 supplemented with 10% of FCS, 1% non-essential amino acids, 1% Hepes, 1% sodium pyruvate and 1% glutamine.

### 2.7 Immunoprecipitation, western blot and *in vitro* protein/protein interaction competition

Cells (5 × 10^6^) were lysed for 20 min at 4°C in lysis buffer (50 mM Tris pH8, 1% NP40, 137 mM NaCl, 1 mM MgCl_2_, 1 mM CaCl_2_, 10% glycerol and protease inhibitor mixture). Lysates (500 μg) were immunoprecipitated with the appropriated antibody overnight at 4°C and protein A/G sepharose was added for 1h at 4°C. After washing with 1x TBST (20 mM Tris-HCl pH7.5, 150 mM NaCl, 0.05% Tween 20), the PP1/LRRK2 interaction was competed using 1 mM of the Mut3DPT-LRRK2-Long peptide for 30 min at room temperature. After several washing steps, immunoprecipitates were separated by SDS-PAGE, transferred to nitrocellulose and blotted with anti PP1 antibody. The membrane was washed and incubated with PO-conjugates secondary antibody. Protein detection was performed using the ECL system. As internal control, the blot was also hybridized with anti-LRRK2 antibody.

### 2.8 Quantification of cellular internalization

Human cell line MDA-MB231 was seeded in 24 well plate (1 × 10^5^ cells/well) and treated with different concentrations of FITC-labelled peptides or for different periods of time. After treatment, cells were harvested and washed twice with PBS to remove the extracellular unbound peptide and resuspended in 200 μL of PBS. FITC fluorescence intensity of internalized peptides was measured by flow cytometry on a FACSCanto II (Beckton Dickinson). Data were analyzed with FACSDiva 6.1.3 software (DB Biosciences). Untreated cells were used as control.

### 2.9 Peptide internalization visualization

For intracellular localization of FITC-labelled peptides, MDA-MB231 cells were seeded in a 8 well Labtek (Thermo Fischer). Cells were treated with FITC-labelled peptides for 4 h and fixed with 4% of formaldehyde for 15 min at room temperature. Samples were washed twice with PBS and mounted in mounting buffer. Images were captured with a fluorescence microscopy (Olympus Japan) using 63x magnification objective.

### 2.10 Characterization of PP1 and LRRK2 peptide interaction by ELISA

A total of 100 μL of biotinylated peptides diluted at 100 μM in PBS were incubated for 2 h at room temperature in a 96-well NeutrAvidin™ coated plate (Pierce, #15128). Wells were washed five times with PBS/0.05% Tween-20 (PBST) and filled with 100 μL of PP1 (Sigma) diluted in PBS/2.5% BSA at the indicated dilutions. Plate were incubated over night at 4°C and washed five times with PBST. Hundred microliters of rabbit polyclonal IgG anti-human PP1α (FL-18) (Santa Cruz Biotechnology, #sc-443) were added at 5 μg/mL in PBS/BSA for 1 hour at room temperature. Wells were washed five times with PBST and filled with 100 μL of HRP conjugated anti-rabbit IgG (Sigma, #A-0545) diluted at 1:20,000 in PBS/BSA for 1 hour at room temperature. Wells were washed five times with PBST and 100 μL of TMB substrate (Pierce, #34021) were added and incubated for 15–45 min. The reaction was stopped with 50 μL of 2 N sulfuric acid, and the absorbance was measured at 450 nm on a Multiskan EX plate reader (Thermo Scientific).

## 3 RESULTS

### 3.1 Identification of LRRK2 sequences involved in PP1 interaction

To determine the amino acid residues of LRRK2 that interact with the serine/threonine phosphatase PP1, we used the PEP-scan approach. Overlapping dodecapeptides covering the whole sequence of LRRK2 protein were immobilized on a cellulose membrane and hybridized with the PP1 protein (Figure 1). A set of 4 contiguous spots revels the presence of a linear interacting area, spanning the residues 1701 to 1718 of the LRRK2 sequence. The domain encompassing this region includes amino acids 1512 to 1878, a region not annotated in the uniprot entry Q5S007. Homology modelling was successful for the region 1688 to 1829, i.e. a domain encompassing the interfering fragment. As depicted in Figure 2A, this LRRK2 fragment corresponds to a region exposed to the solvent, consistently with its interfering ability and adopts an α helical conformation. Such helical conformation is also predicted for the peptide in isolation (Figure 2B) by the PEP-FOLD software. The search for sequence variants was performed using the blastp facility of the Uniprot server for the modelled sequence and searching the Uniprot mammals (resp. vertebrates) sequences, using the default values for the parameters proposed by the Uniprot server, and selecting only the sequences with an e-value less than 0.00001. Over the 129 (resp. 281) matches identified on date of 2019 June 11, it revealed a strong conservation at most positions of the interfering fragment (Figure 2C), 8 (resp. 6) positions upon 18 being strictly conserved among the mammal sequences (resp. vertebrates), and most other positions showing only limited variations, suggesting possible functional constraints for this fragment.

**Figure 1.**
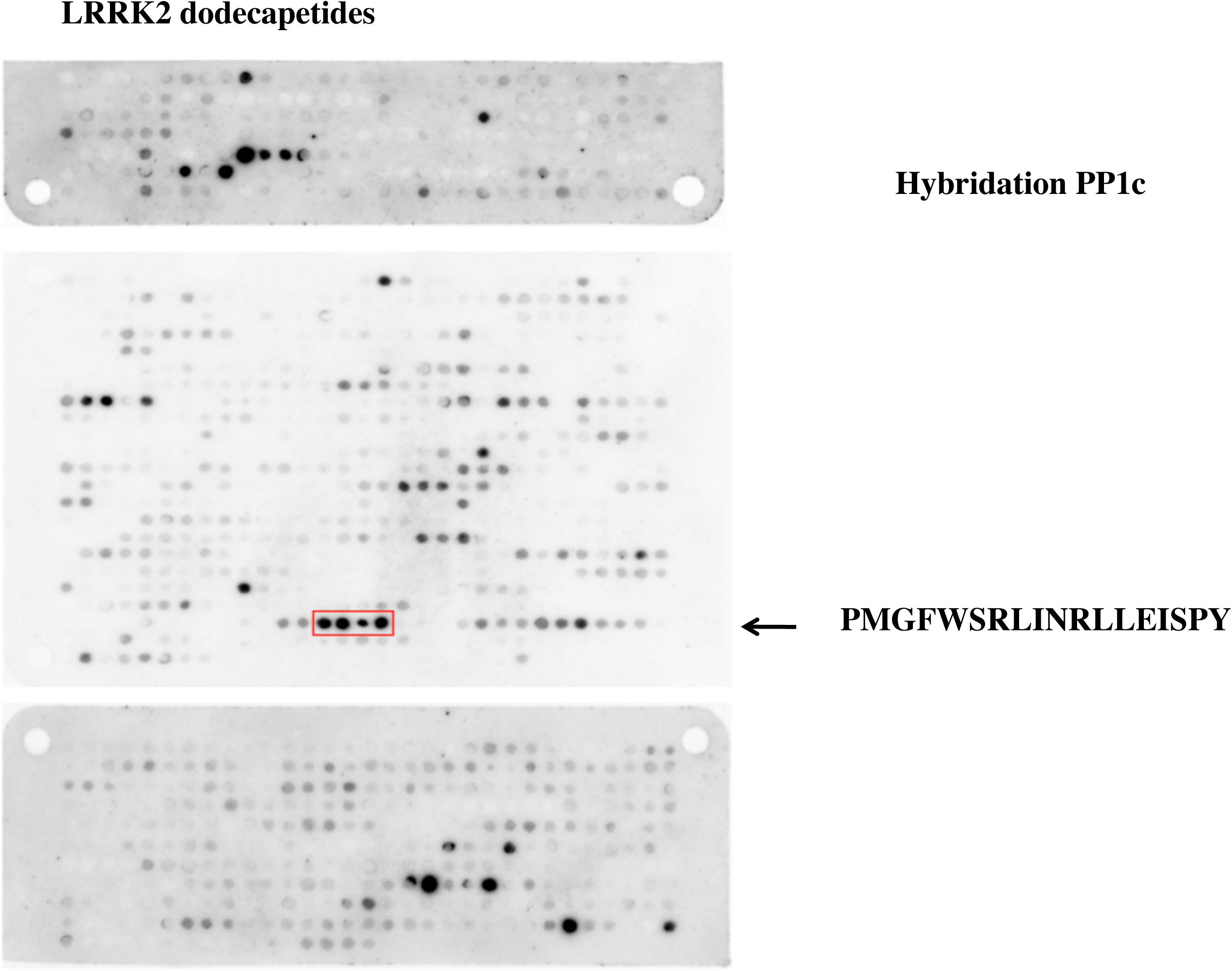
Identification of the binding site of LRRK2 to PP1c. The sequence of LRRK2 was developed as series of overlapping dodecapeptides with a shift of two amino acids. The membrane was hybridized with purified PP1c protein, followed by a secondary antibody. Spots were detected using the ECL system. LRRK2 peptides that interact with PP1c are boxed and the sequence shown.

**Figure 2.**
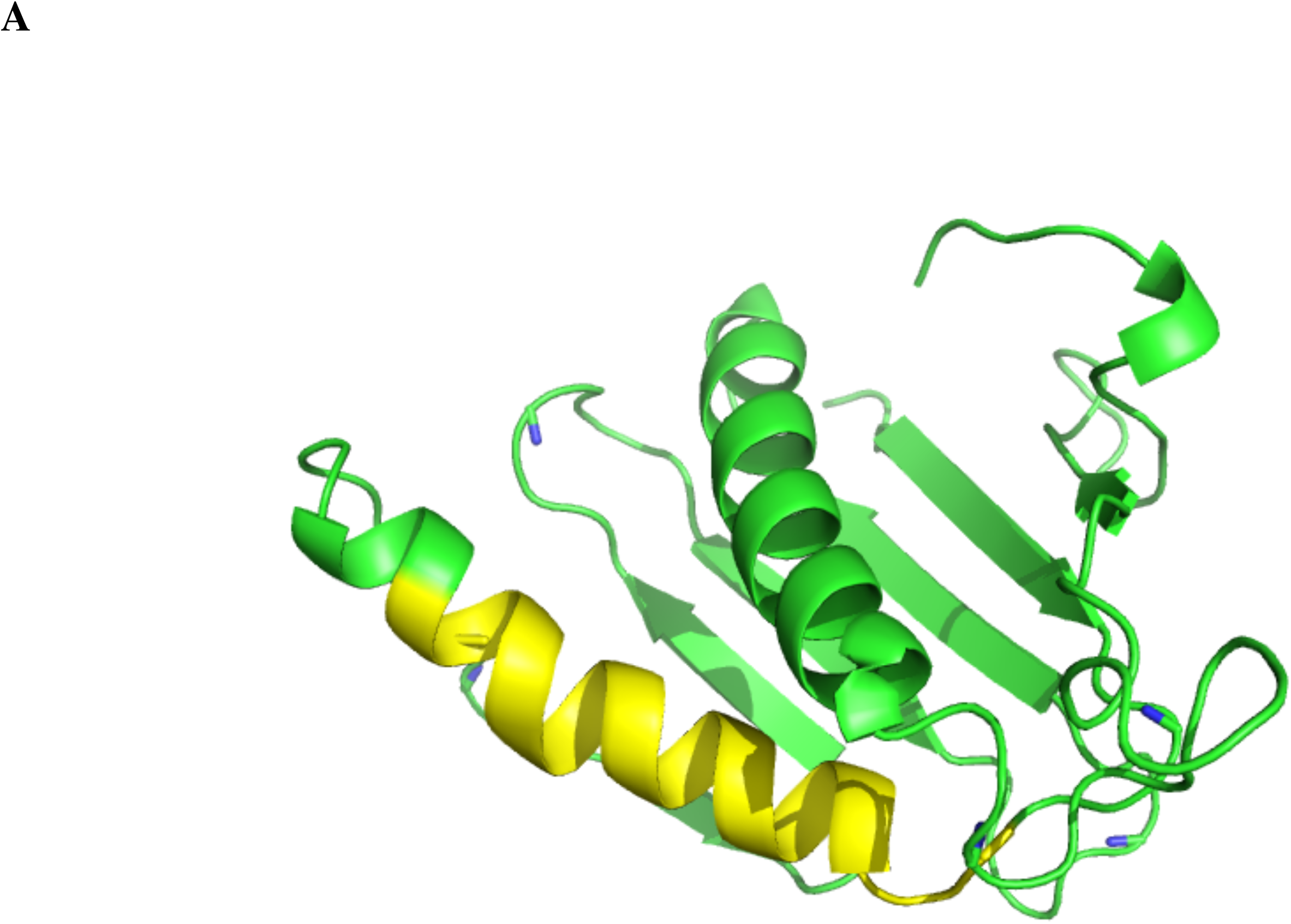
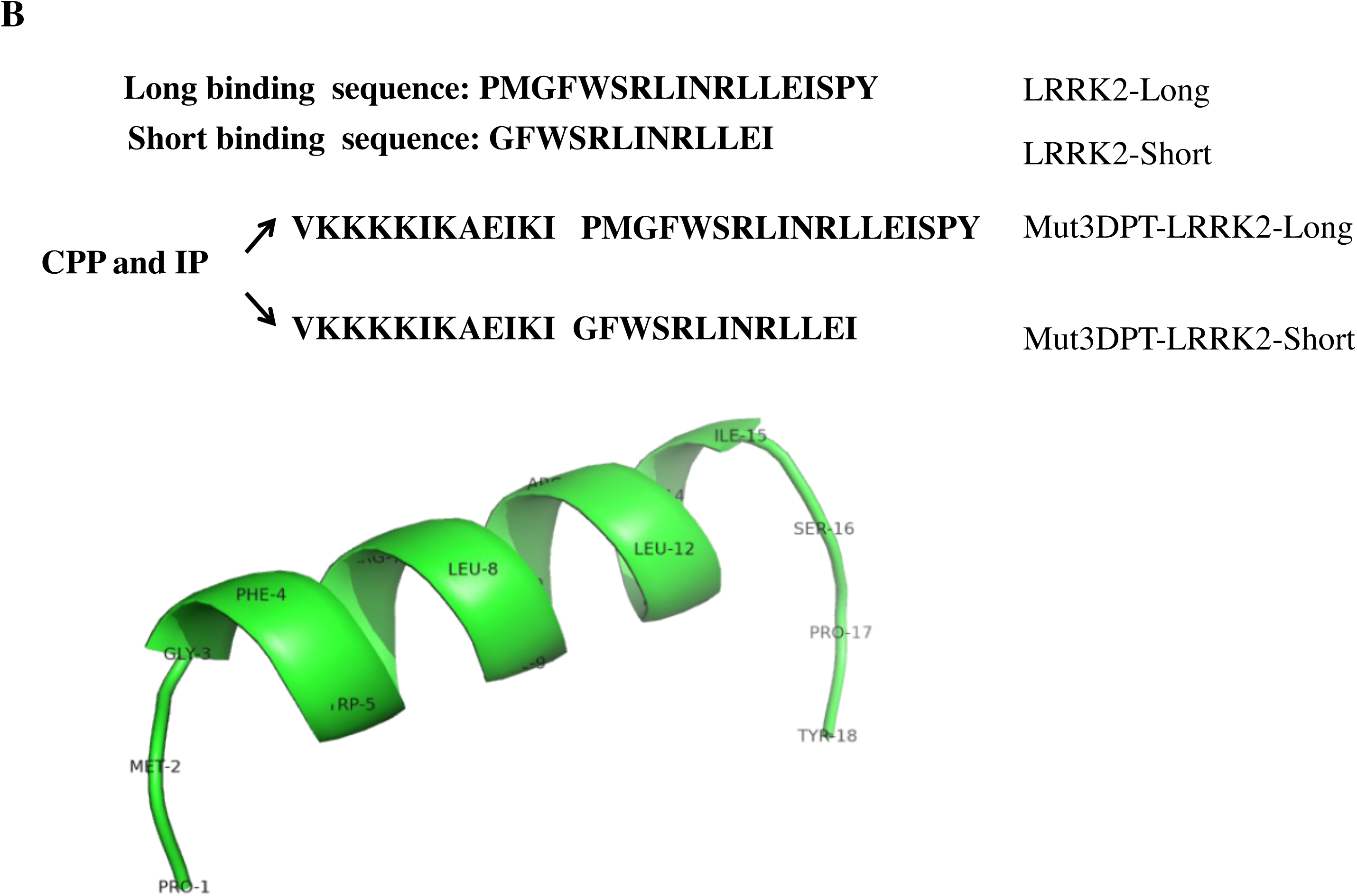
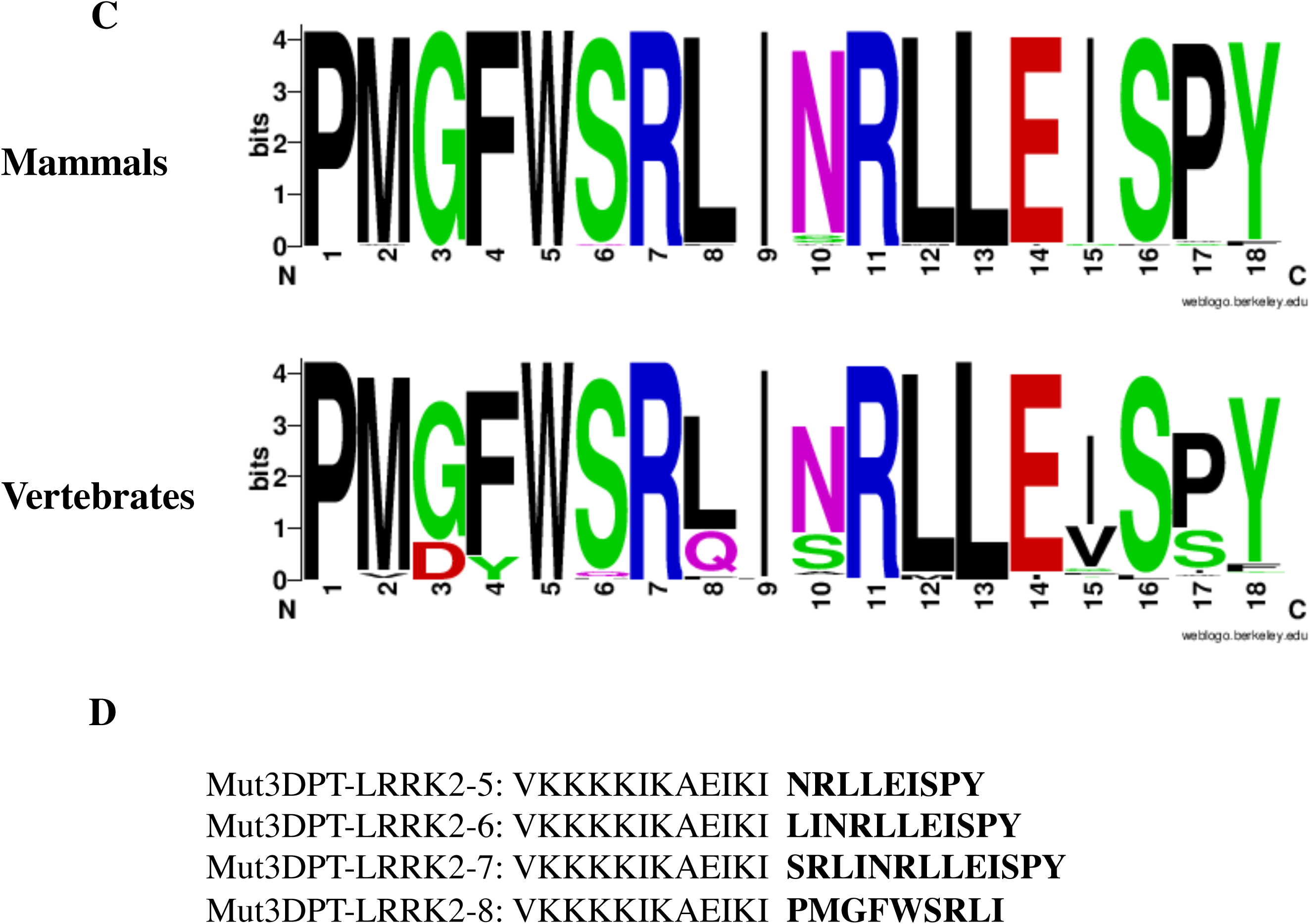
Structure of the Short and Long sequence of the LRRK2 interacting peptides. A) Homology model of the LRRK2 domain encompassing the interfering fragment (depicted in yellow). B) Sequence of the short and long interfering peptides. The sequences were associated to a cell penetrating peptide named Mut3DPT, previously described for generation of the cell penetrating and interfering peptides (CP and IP). The helical structure of the long interfering peptide is also shown, indicating the amino acids that were deleted in the short version of the peptide. C) Sequence variation observed through related Uniprot mammal (top) and vertebrate (bottom) sequences. D) Sequences of the new generated versions of the Mut3DPT-LRRK2-Long peptide, LRRK2-5 to LRRK2-8.

Chimeric peptide containing Mut3DPT-Sh1 (VKKKKIKAEIKI), an optimized cell penetrating peptide, fused to the newly identified interaction sequence of LRRK2 to PP1 was chemically synthesized. This peptide, named Mut3DPT-LRRK2-Long, was further used for functional analysis. We then analysed whether the original interaction sequence could be shortened. As shown on Figure 2B, the peptide is predicted to adopt a helical conformation, but its N- and C-terminal regions seem less structured. In order to possibly minimize the entropic cost upon peptide binding, a shorter peptide was designed by removing the 2 first and the 3 last amino acids of the long peptide (Mut3DPT-LRRK2-Short).

The interesting results obtained with both peptides promotes us to check whether the whole sequence of Mut3DPT-LRRK2-Long peptide was indispensable for blocking the interaction LRRK2/PP1, Using PEP-FOLD prediction software, shorter versions of the Mut3DPT-LRRK2-Long peptide were designed with the aim to identify the short versions of Mut3DPT-LRRK2-Long able to bind PP1. According to this, four new peptides were generated (LRRK2-5, LRRK2-6, LRRK2-7 and LRRK2-8) and associated to the shuttle Mut3DPT (Fig 2D).

### *3.2 In vitro* competition of LRRK2/PP1 interaction

An *in vitro* competition assay was done in order to confirm that Mut3DPT-LRRK2-Long peptide specifically targets the LRRK2/PP1 interaction. Lysates from MDA-MB231 cell line were immunoprecipitated with anti-LRRK2 antibody and the interaction with PP1 was competed using Mut3DPT-LRRK2-Long peptide (Figure 3A). PP1 protein was detected in control LRRK2 immunoprecipitates and in immunoprecipitates competed using the shuttle alone (Mut3DPT), while it was low detected after competition with 1 mM of Mut3DPT-LRRK2-Long peptide. LRRK2 was used as internal control showing similar level in all conditions. This result suggests that Mut3DPT-LRRK2-Long peptide specifically target the interaction between human LRRK2 and PP1.

**Figure 3.**
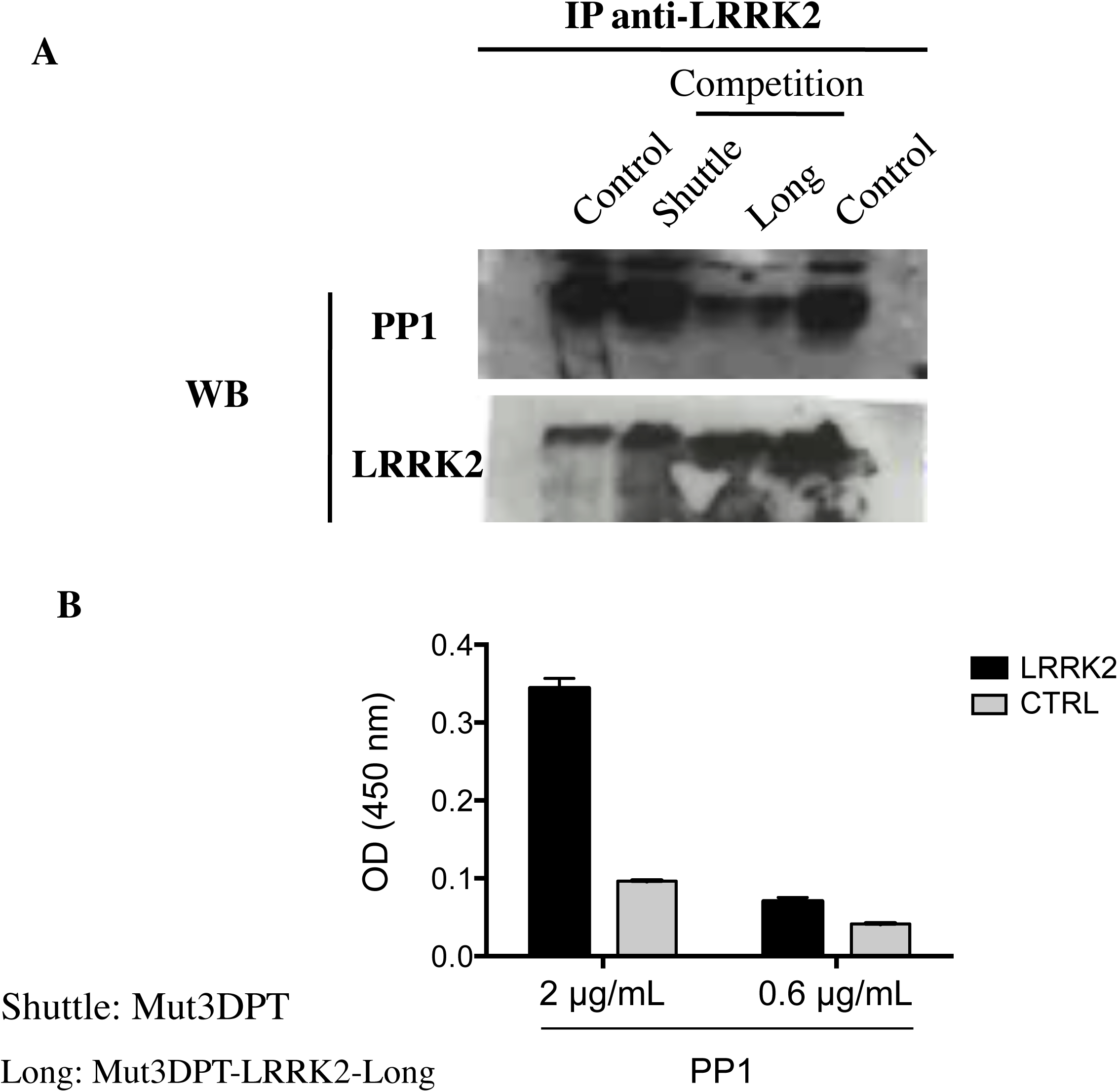
Mut3-DPT-LRRK2-Long peptide competes *in vitro* LRRK2/PP1c interaction. A) The MDA-MB231 cell line was lysed and cytoplasmic extracts immunoprecipitated with anti-LRRK2 antibody. The LRRK2/PP1c interaction was competed *in vitro* with 1 mM of Mut3DPT-LRRK2-Long peptide for 30 min at room temperature. Immunoprecipitates were washed and immunoblotted with anti-PP1c and anti-LRRK2 antibody, the former as internal control of protein loading. B) Mut3DPT-LRRK2-Long and control (CTRL) biotinylated peptides were immobilized on a NeutrAvidin™ coated plate and incubated overnight with dilutions of PP1α at 2 and 0.6 μg/mL. After washing, rabbit anti-PP1α was added in each well and incubated 1 hour at room temperature. Wells were washed and filled with a dilution of HRP-conjugated anti-Rabbit. Binding activity of PP1α is expressed as mean OD at 450 nm of duplicate wells, and bars indicate SD. These data are representative of two independent experiments.

Then, we analysed whether the biotinylated Mut3DPT-LRRK2-Long peptide was able to associate with purified PP1 protein. Figure 3B shows that the peptide recognizes the protein PP1 in an ELISA test. An irrelevant peptide was used as negative control.

In the same way, we investigated whether the shortest versions of the original peptide, Mut3DPT-LRRK2-5 to Mut3DPT-LRRK2-8, were able to compete with the LRRK2/PP1 interaction. Figure 4A shows that none of the shortest versions of the original peptide were able to compete *in vitro* the interaction LRRK2/PP1 using them at concentration of 1 mM, as in figure 3A. As positive control, the interaction was competed in anti-LRRK2 immunoprecipitates using 1 mM of the original Mut3-DPT-LRRK2-Long peptide (Fig 4A). To confirm this result, we analysed whether the peptides Mut3DPT-LRRK2-5 to Mut3DPT-LRRK2-8 were able to associate to purified PP1 protein in an ELISA test using biotinylated peptides. Figure 4B shows that none of the peptides recognize purified PP1 protein, confirming the absence of in vitro competition of the interaction.

**Figure 4.**
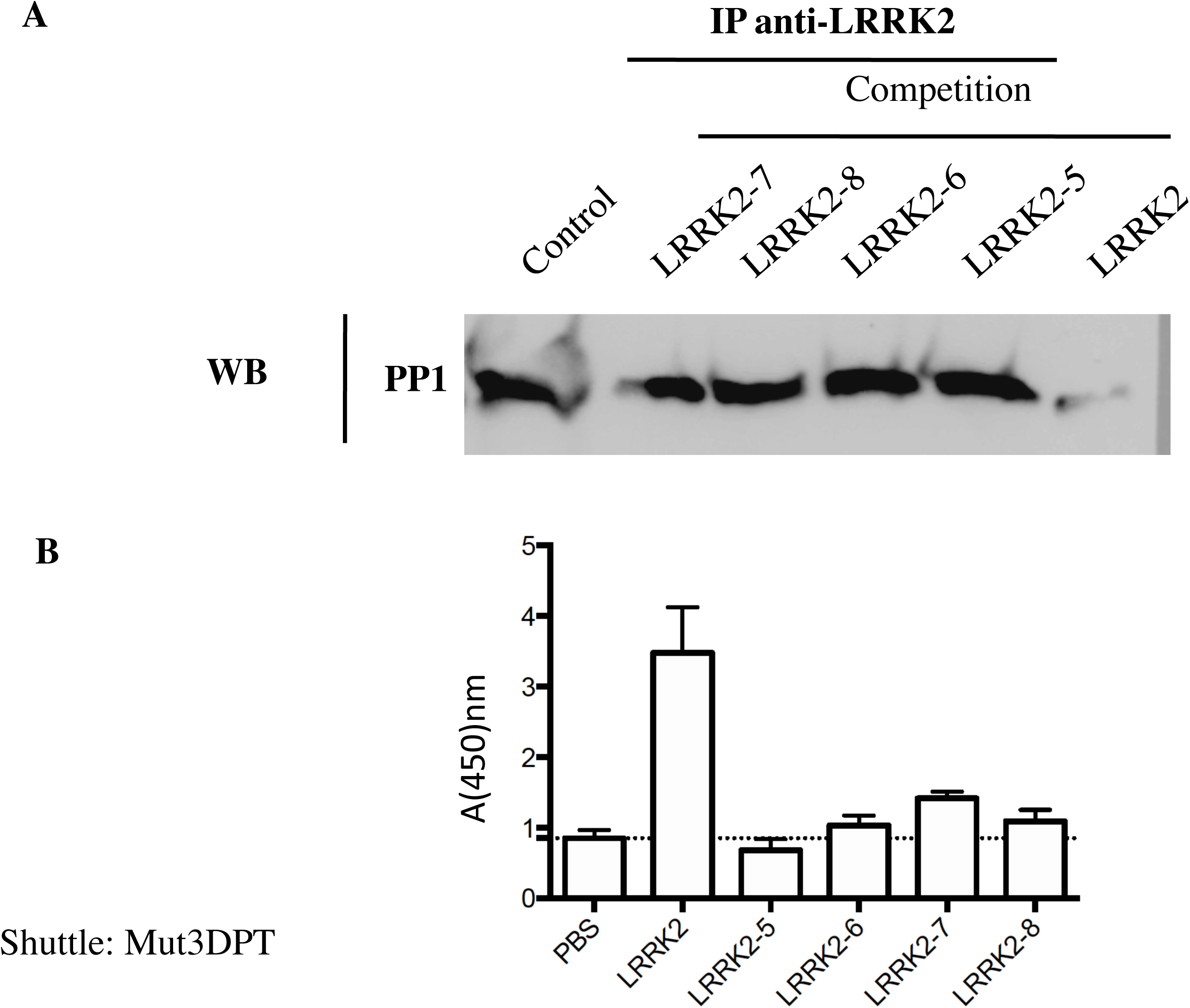
*In vitro* competition of LRRK2/PP1c interaction using LRRK2-5 to LRRK2-8 peptides. A) The MDA-MB231 cell line was lysed and cytoplasmic extracts immunoprecipitated with anti-LRRK2 antibody. The LRRK2/PP1c interaction was competed *in vitro* with 1 mM of peptide for 30 min at room temperature. Immunoprecipitates were washed and immunoblotted with anti-PP2c. B) Biotinylated peptides were immobilized on a NeutrAvidin™ coated plate and incubated overnight with purified PP1α protein. After washing, rabbit anti-PP1α was added in each well and incubated 1 hour at room temperature. Wells were washed and filled with a dilution of HRP-conjugated anti-Rabbit. Binding activity of PP1α is expressed as mean OD at 450 nm of duplicate wells, and bars indicate SD.

Taken together, we can conclude that the active peptides that recognize PP1 are Mut3-DPT-LRRK2-Long and Mut3DPT-LRRK2-Short. The other variants lost the capacity to recognize the protein PP1.

### 3.3 Quantification of internalization of Short and Long version of interfering peptides

We evaluated whether the peptides Mut3DPT-LRRK2-Long and Mut3DPT-LRRK2-Short peptides were able to internalize into cells. The peptides were labelled with FITC and internalization was analysed by FACS. MDA-MB231 cells were treated with FITC-labelled Mut3DPT-LRRK2-Long and Mut3DPT-LRRK2-Short peptides at different concentrations for 4h and then, internalization analysed by FACS. Figure 4A shows that the fluorescence intensity detected is higher when using the Mut3DPT-LRRK2-Short peptide, compared to Mut3DPT-LRRK2-Long peptide, which shows lower fluorescence intensity. We next analysed the effect of time incubation in the internalization at a fixed concentration of peptides. Again, the Mut3DPT-LRRK2-Short peptide shows higher level of internalization, reaching a plateau upon 3 h of incubation (Figure 5B). Taken together, our results suggest that Mut3DPT-LRRK2-Short peptide shows more favourable internalization parameters that Mut3DPT-LRRK2-Long peptide.

**Figure 5.**
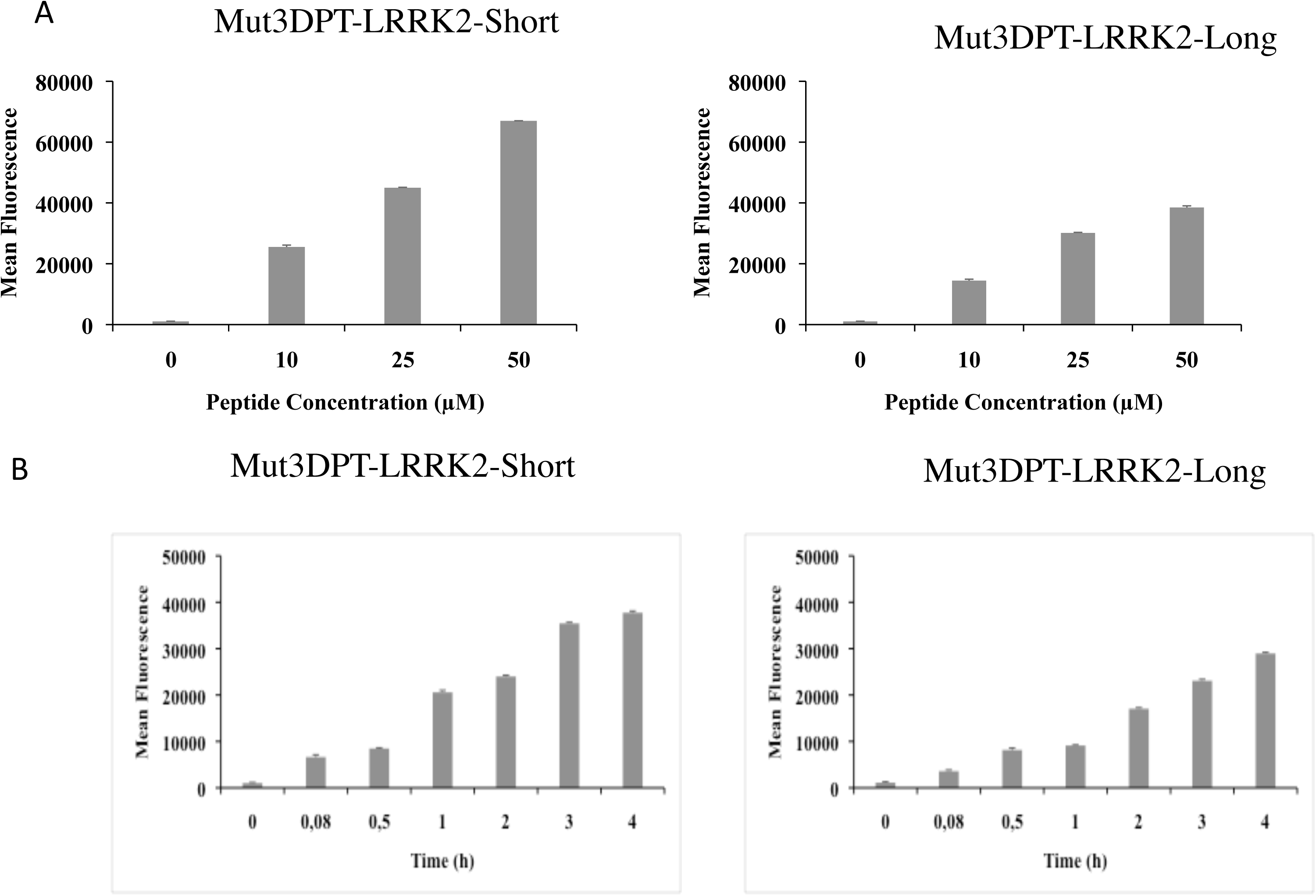
Concentration and time-dependent internalization of FITC-labelled Mut3DPT-LRRK2-Short and Mut3DPT-LRRK2-Long peptides. A) MDA-MB231 cells were incubated 4h with different concentrations of FITC-labelled peptides. The mean fluorescence intensity was detected by flow cytometry and compared to control non-treated control cells. Standard deviation is shown. B) MDA-MB231 cells were incubated with 20 μM of the FITC-labelled peptides for different periods of time. The mean fluorescence intensity was detected by flow cytometry. Non-treated cells were used as control. Bars indicate standard deviation.

### 3.4 Internalization of Mut3DPT-LRRK2-Long and Mut3DPT-LRRK2-Short peptides into primary cells

In addition to cell lines, we also tested the internalization of the Mut3DPT-LRRK2-Short and Mut3DPT-LRRK2-Long in peripheral blood mononuclear cells (PBMC) from healthy donors and chronic lymphocytic leukemia (CLL) patients. PBMC from both were incubated with 50 μM of both peptides for 4h at 37°C. As illustrated in Figure 6, the Mut3DPT-LRRK2-Short peptide shows slightly higher fluorescence intensity than the Mut3DPT-LRRK2-Long peptide in healthy donors and CLL patients. This result is in accordance to those obtained using cell line MDA-MB231 and confirms also that, usually, shorter peptides are better internalized.

**Figure 6.**
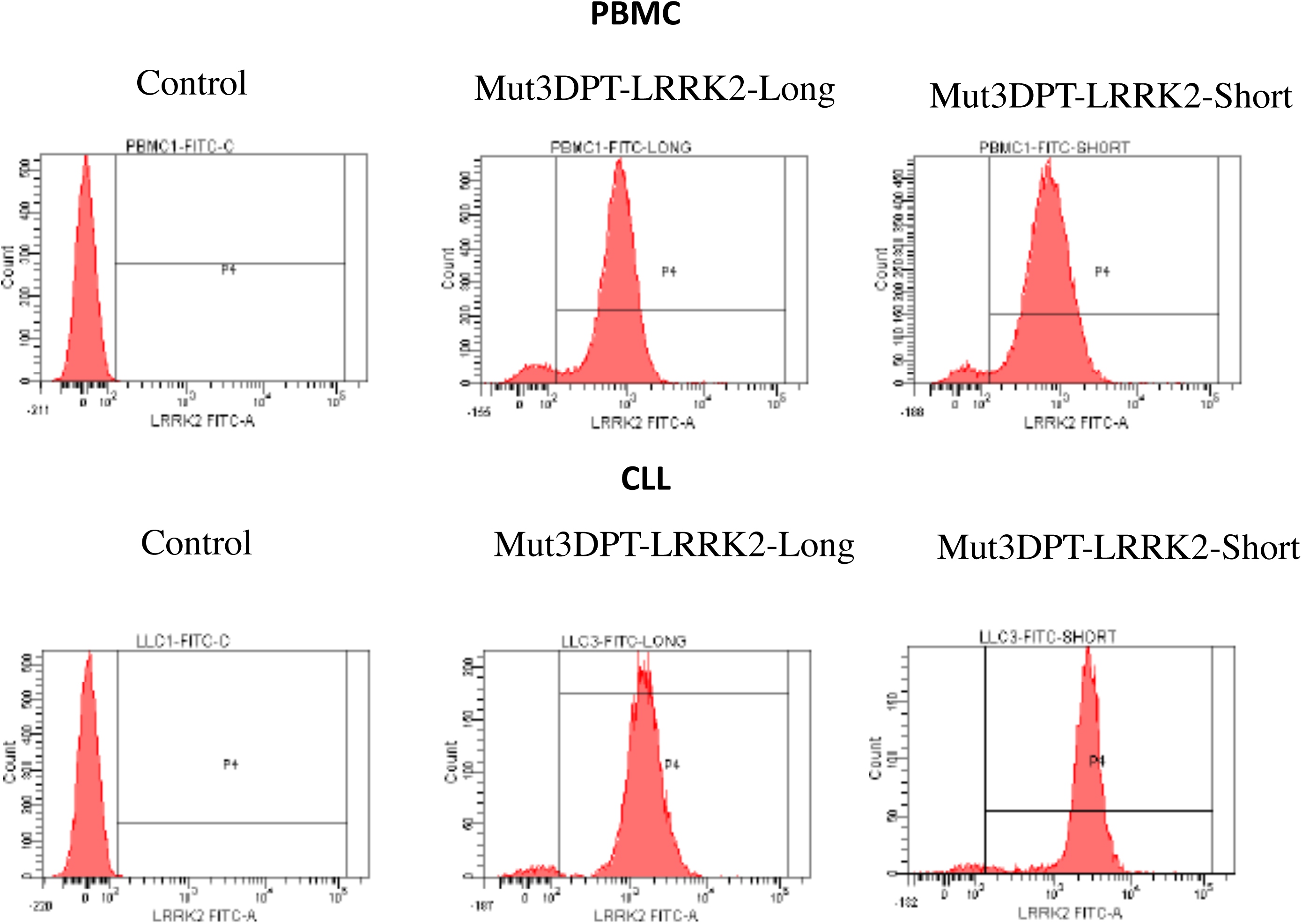
Internalization of FITC-labelled Mut3DPT-LRRK2-Short and Mut3DPT-LRRK2-Long peptides on healthy and tumoral PBMC. Peripheral blood mononuclear cells (PBMC) of healthy donors (HD) or chronic lymphocytic leukemia (CLL) patients were incubated 4h with 50 μM of the FITC-labelled Mut3DPT-LRRK2-Short and Mut3DPT-LRRK2-Long peptides. The mean fluorescence intensity was detected by flow cytometry. Non-treated cells were used as control.

### 3.5 Intracellular localization of the Mut3DPT-LRRK2-Short and Mut3DPT-LRRK2-Long peptides

To confirm the FACS results and to visualize the intracellular distribution of Mut3DPT-LRRK2-Short and Mut3DPT-LRRK2-Long peptides, we stained MDA-MB231 cells with FITC-labelled peptides and analysed the distribution by fluorescence microscopy following 2h of incubation at a concentration of 25 μM (Figure 7). The staining pattern of both peptides is similar with a punctuate staining that could maybe corresponds to association to a given organelle. Again, Mut3DPT-LRRK2-Long peptide shows slightly weak fluorescence intensity into the cells.

**Figure 7.**
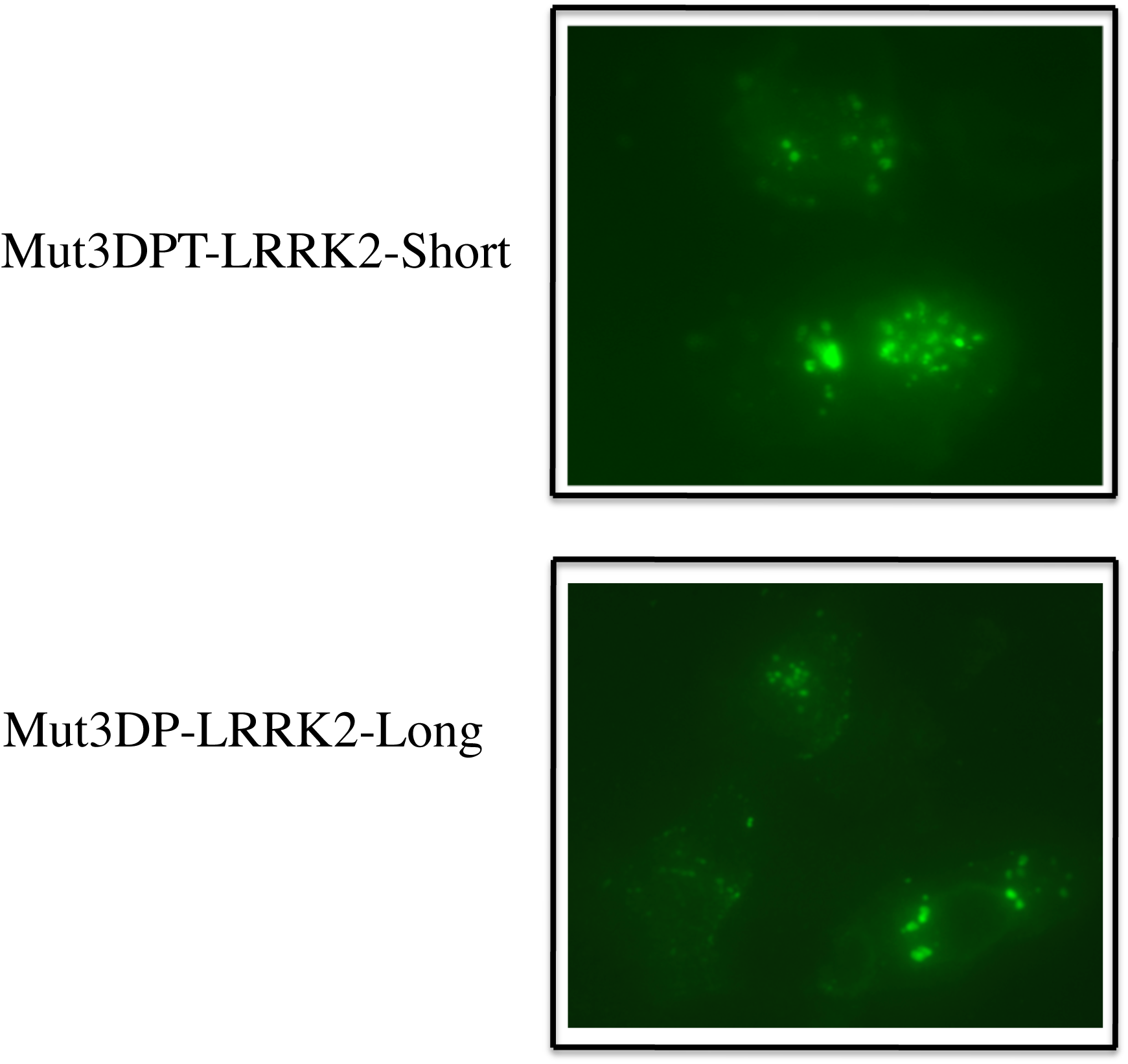
Intracellular localization of FITC-labelled Mut3DPT-LRRK2-Short and Mut3DPT-LRRK2-Long peptides. MDA-MB231 cells were grown on coverslips and incubated 1h at 37°C with 25 μM of FITC-labelled Mut3DPT-LRRK2-Short and Mut3DPT-LRRK2-Long peptides. Cells were washed 3 times with PBS, fixed with 4% paraformaldehyde (PFA) and analysed by fluorescence microscopy.

## 4 DISCUSSION

Several serine/threonine protein kinase modulators (particularity inhibitors) have been developed and some of them have reached therapeutically developments. Nevertheless, a very few number of serine/threonine protein phosphatase modulators are available. In parallel, protein-protein interactions (PPI) regulate subtly and through different ways cell homeostasis. Consequently, deregulation of these interactions that are often associated with pathology, could be a good opportunity to understand PPI effects ^23^. For these reasons, we are interested in developing peptides acting as phosphatases modulators by targeting the enzymatic or the partners’ interacting regions. We have developed interfering peptides (IPs) targeting interaction of PP2 with some of its partners ^14^. In this manuscript, we have targeted the interaction between PP1 and LRRK2, a Parkinson disease associated protein. As PP1 is a hub protein, we decided that it would be best to identify peptides targeting PP1 with LRRK2 derived peptides, because chances to get unpredicted effect using PP1-derived peptides seem be high. The sequence of LRRK2 interacting with PP1 contains eighteen amino acids (PMGFWSRLINRLLEISPY) forming an alpha helix structure and located in an exposed region of the LRRK2 protein. Starting from this sequence and using *in silico* approaches, five new IPs were designed. These sequences were fused to an optimized penetrating peptide ^13^ to generate chimeric peptides able to penetrate into the cell and to block the interaction PP1/LRRK2. The specific interfering activity of these peptides was confirmed using *in vitro* competition assays. As the Mut3DPT-LRRK2-Long and Mut3DPT-LRRK2-Short peptides were fused to an immune fluorescent marker, we were able to show that the shorter version of the peptide has a better internalization either in cells lines or in primary cells.

Mut3DPT-LRRK2-Long displays the best performance in terms of PP1/LRRK2 modulation, compared to results obtained for Mut3DPT-LRRK2-Short and Mut3DPT-LRRK2-5- to 8. The poorer performance of Mut3DPT-LRRK2-Short compared to Mut3DPT-LRRK2-Long suggests the importance of the residues located ant N-ter and C-ter extremities of Mut3DPT-LRRK2-Long. The inactivity of Mut3DPT-LRRK2-5- to 8, which include those residues, suggests that either several hot spots dispatched along Mut3DPT-LRRK2-Long are necessary to drive the binding to PP1, or that the structural properties of the half-peptides are not those expected (off-target and non specific binding could occur), or the peptide conformation could be less stable than expected, possibly when linked to Mut3DPT.

The LRRK2 protein is phosphorylated at multiple sites but the regulation of its phosphorylation is not fully understood ^10,24-27^. Changes in the LRRK2 phosphorylation status are linked to the pathogenesis of LRRK2-related PD and the available data show that phosphorylation is a highly regulated physiological event in the disease, It has been shown that PP1 is the phosphatase that efficiently dephosphorylates LRRK2 ^8^, although there is also evidences that other serine/threonine phosphatase, like PP2A, may play an auxiliary role in dephosphorylation of LRRK2 under specific conditions. It has been published that LRRK2 dephosphorylation involves the enhanced access of PP1 to LRRK2 phosphosites. It has been also shown that most of LRRK2 PD mutations have decreased phosphorylation at several Ser residues ^24,25,27^. According to this, our cell penetrating and interfering peptide is an important tool to control PP1/LRRK2 association and, as a consequence, the phosphorylation of LRRK2. Taken together, our manuscript proposes new tools to manipulate and to study the PP1/LRRK2 interaction in normal and pathological conditions, such as PD.

## AKDNOWLEDGEMENTS

This work was supported by Inserm and Paris Diderot University.

## REFERENCES

1 Kolupaeva, V. & Janssens, V. PP1 and PP2A phosphatases--cooperating partners in modulating retinoblastoma protein activation. The FEBS journal 280, 627–643, doi:10.1111/j.1742-4658.2012.08511.x (2013).

2 Takai, A. et al. Protein phosphatases 1 and 2A and their naturally occurring inhibitors: current topics in smooth muscle physiology and chemical biology. The journal of physiological sciences: JPS 68, 1–17, doi:10.1007/s12576-017-0556-6 (2018).

3 Verbinnen, I., Ferreira, M. & Bollen, M. Biogenesis and activity regulation of protein phosphatase 1. Biochemical Society transactions 45, 89–99, doi:10.1042/BST20160154 (2017).

4 Garcia, A. et al. Serine/threonine protein phosphatases PP1 and PP2A are key players in apoptosis. Biochimie 85, 721–726 (2003).

5 Heroes, E. et al. The PP1 binding code: a molecular-lego strategy that governs specificity. The FEBS journal 280, 584–595, doi:10.1111/j.1742-4658.2012.08547.x (2013).

6 Sheppeck, J. E., 2nd, Gauss, C. M. & Chamberlin, A. R. Inhibition of the Ser-Thr phosphatases PP1 and PP2A by naturally occurring toxins. Bioorganic & medicinal chemistry 5, 1739–1750 (1997).

7 Reynhout, S. & Janssens, V. Physiologic functions of PP2A: Lessons from genetically modified mice. Biochimica et biophysica acta. Molecular cell research 1866, 31–50, doi:10.1016/j.bbamcr.2018.07.010 (2019).

8 Lobbestael, E. et al. Identification of protein phosphatase 1 as a regulator of the LRRK2 phosphorylation cycle. The Biochemical journal 456, 119–128, doi:10.1042/BJ20121772 (2013).

9 Taymans, J. M. & Baekelandt, V. Phosphatases of alpha-synuclein, LRRK2, and tau: important players in the phosphorylation-dependent pathology of Parkinsonism. Frontiers in genetics 5, 382, doi:10.3389/fgene.2014.00382 (2014).

10 Lobbestael, E., Baekelandt, V. & Taymans, J. M. Phosphorylation of LRRK2: from kinase to substrate. Biochemical Society transactions 40, 1102–1110, doi:10.1042/BST20120128 (2012).

11 Inzelberg, R. et al. The LRRK2 G2019S mutation is associated with Parkinson disease and concomitant non-skin cancers. Neurology 78, 781–786, doi:10.1212/WNL.0b013e318249f673 (2012).

12 Meeusen, B. & Janssens, V. Tumor suppressive protein phosphatases in human cancer: Emerging targets for therapeutic intervention and tumor stratification. The international journal of biochemistry & cell biology 96, 98–134, doi:10.1016/j.biocel.2017.10.002 (2018).

13 Zhang, X., Brossas, J. Y., Parizot, C., Zini, J. M. & Rebollo, A. Identification and characterization of novel enhanced cell penetrating peptides for anti-cancer cargo delivery. Oncotarget 9, 5944–5957, doi:10.18632/oncotarget.23179 (2018).

14 Arrouss, I. et al. Specific targeting of caspase-9/PP2A interaction as potential new anti-cancer therapy. PloS one 8, e60816, doi:10.1371/journal.pone.0060816 (2013).

15 Dominguez-Berrocal, L. et al. New Therapeutic Approach for Targeting Hippo Signalling Pathway. Scientific reports 9, 4771, doi:10.1038/s41598-019-41404-w (2019).

16 Lamiable, A. et al. PEP-FOLD3: faster de novo structure prediction for linear peptides in solution and in complex. Nucleic acids research 44, W449–454, doi:10.1093/nar/gkw329 (2016).

17 Frank, R. & Overwin, H. SPOT synthesis. Epitope analysis with arrays of synthetic peptides prepared on cellulose membranes. Methods in molecular biology 66, 149–169, doi:10.1385/0-89603-375-9:149 (1996).

18 Gausepohl, H., Boulin, C., Kraft, M. & Frank, R. W. Automated multiple peptide synthesis. Peptide research 5, 315–320 (1992).

19 Soding, J. Protein homology detection by HMM-HMM comparison. Bioinformatics 21, 951–960, doi:10.1093/bioinformatics/bti125 (2005).

20 Berman, H. M. et al. The Protein Data Bank. Nucleic acids research 28, 235–242, doi:10.1093/nar/28.1.235 (2000).

21 Catherinot, V. & Labesse, G. ViTO: tool for refinement of protein sequence-structure alignments. Bioinformatics 20, 3694–3696, doi:10.1093/bioinformatics/bth429 (2004).

22 Karami, Y., Guyon, F., De Vries, S. & Tuffery, P. DaReUS-Loop: accurate loop modeling using fragments from remote or unrelated proteins. Scientific reports 8, 13673, doi:10.1038/s41598-018-32079-w (2018).

23 Modell, A. E., Blosser, S. L. & Arora, P. S. Systematic Targeting of Protein-Protein Interactions. Trends in pharmacological sciences 37, 702–713, doi:10.1016/j.tips.2016.05.008 (2016).

24 Doggett, E. A., Zhao, J., Mork, C. N., Hu, D. & Nichols, R. J. Phosphorylation of LRRK2 serines 955 and 973 is disrupted by Parkinson’s disease mutations and LRRK2 pharmacological inhibition. Journal of neurochemistry 120, 37–45, doi:10.1111/j.1471-4159.2011.07537.x (2012).

25 Dzamko, N. et al. Inhibition of LRRK2 kinase activity leads to dephosphorylation of Ser(910)/Ser(935), disruption of 14-3-3 binding and altered cytoplasmic localization. The Biochemical journal 430, 405–413, doi:10.1042/BJ20100784 (2010).

26 Li, X. et al. Phosphorylation-dependent 14-3-3 binding to LRRK2 is impaired by common mutations of familial Parkinson’s disease. PloS one 6, e17153, doi:10.1371/journal.pone.0017153 (2011).

27 Nichols, R. J. et al. 14-3-3 binding to LRRK2 is disrupted by multiple Parkinson’s disease-associated mutations and regulates cytoplasmic localization. The Biochemical journal 430, 393–404, doi:10.1042/BJ20100483 (2010).

